## Supplementary Information for "A microfluidic device for inferring metabolic landscapes in yeast monolayer colonies"

### Supplementary Table 1

| Name | Background | Genotype | Source |
|---|---|---|---|
| **yPH001** | BY4741 | MATa his3Δ1 leu2Δ0 met15Δ0 ura3Δ0 | Leon's Lab (IJM/ CNRS) |
| **yPH152** | BY4741 | HXT1-GFP :HisMX | Leon's Lab (IJM/ CNRS) |
| **yPH155** | BY4741 | HXT7-GFP::HphNT | Leon's Lab (IJM/ CNRS) |
| **yPH179** | BY4741 | HXT2-GFP::HisMX | Yeast GFP collection |
| **yPH180** | BY4741 | HXT3-GFP::HisMX | Yeast GFP collection |
| **yPH182** | BY4741 | GLK1-GFP::HisMX | Yeast GFP collection |
| **yPH183** | BY4741 | MIG1-GFP::HisMX | Yeast GFP collection |
| **yPH188** | BY4741 | PDC1-GFP::HisMX | Yeast GFP collection |
| **yPH189** | BY4741 | HXK1-GFP::HisMX | Yeast GFP collection |
| **yPH190** | BY4741 | HXK2-GFP::HisMX | Yeast GFP collection |
| **yPH191** | BY4741 | SDH2-GFP::HisMX | Yeast GFP collection |
| **yPH192** | BY4741 | HXT4-GFP::HisMX | Leon's Lab (IJM/ CNRS) |
| **yPH193** | BY4741 | HXT5-GFP::HisMX | Leon's Lab (IJM/ CNRS) |
| **yPH236** | BY4741 | HXT6-GFP::Hyg | Leon's Lab (IJM/ CNRS) |

**Supplementary Table 1.** List of yeast *S. cerevisiae* strains used in this study.

### Supplementary Figure 1

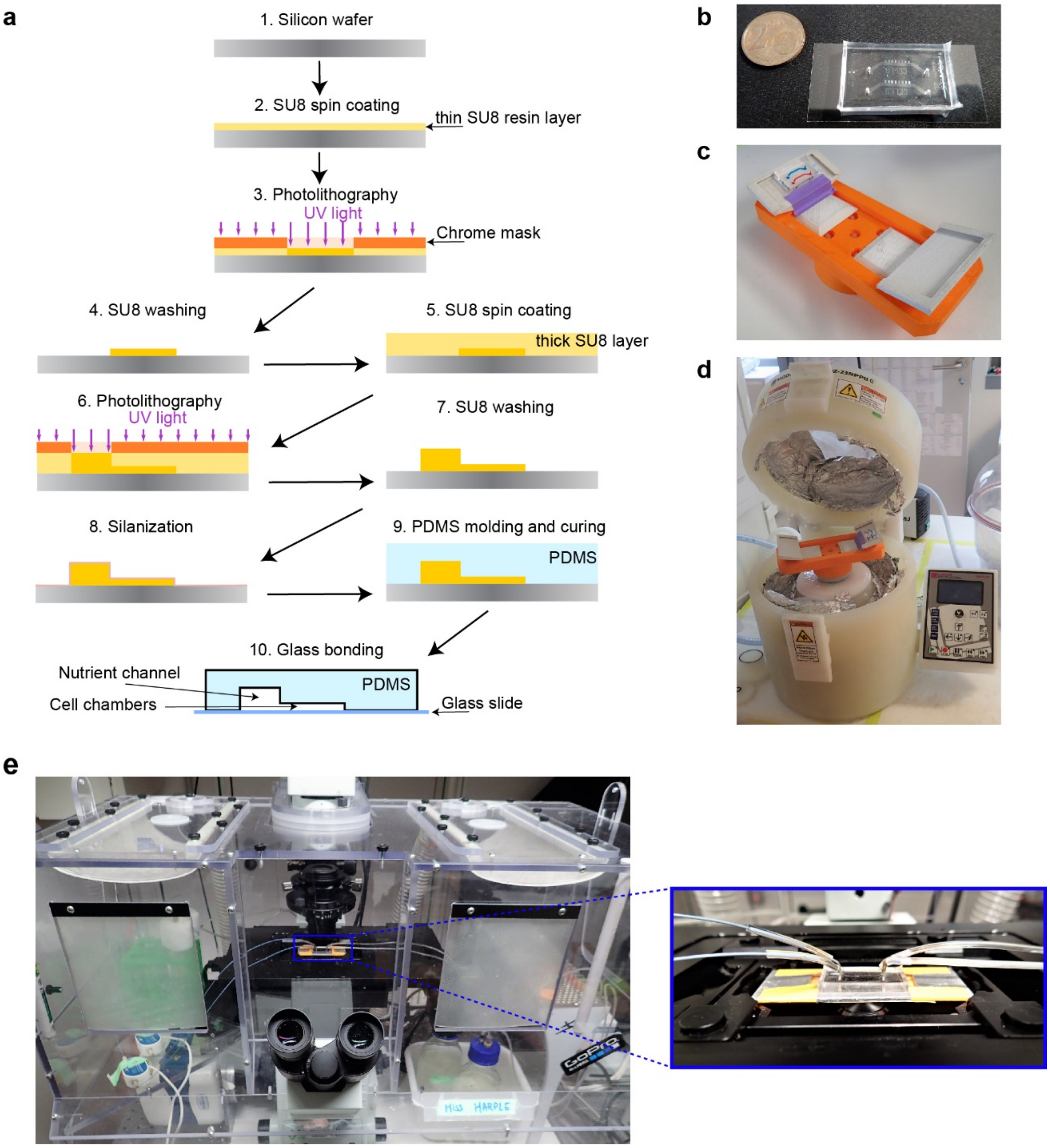

**Supplementary Figure 1. Experimental Details. S1a**. Microfluidic device fabrication. Epoxy-based resin is spread on a silicon wafer and illuminated with the UV light through a custom designed chrome mask. Soluble part of the resin is washed off and a new cycle of resin deposition can be made depending on the number of layers of the microfluidic device. The master wafer is silanized and used as a re-usable negative mold to produce microfluidic chips. Polydimethylsiloxane (PDMS) is then poured on the master wafer and cured until it is polymerized. The PDMS chip is bonded with a glass coverslip by plasma bonding. The final microfluidic chip is used to grow cells and deliver nutrients. The thinner parts are the cell chambers, and the thicker part is the main nutrient channel. **S1b.** Microfluidic chip bonded to a cover slip. Each chip has two independent "yeast machines". **S1c.** 3D printed chip holder for centrifugation with the chip mounted on it (channels are colored with different dyes for better visualization). **S1d.** Spin coater that is used for centrifugation of cells into the cell chamber. **S1e.** Microfluidic chip mounted on a microscope.

### Supplementary Figure 2

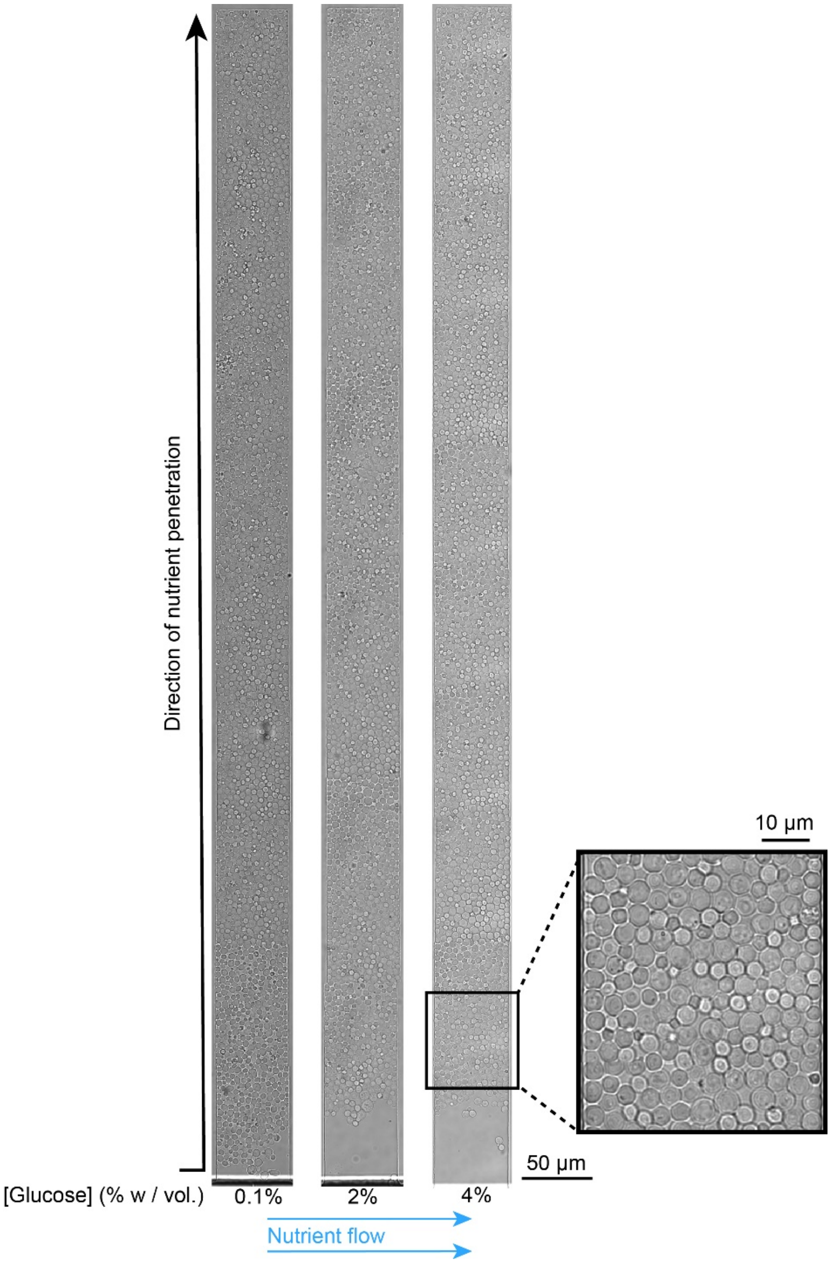

**Supplementary Figure 2. Detailed cell chamber view.** Cell chambers of the "yeast machine" at high magnification in different starting glucose concentrations. Cells are vertically constrained to facilitate single-cell imaging and time-lapse fluorescence microscopy.

### Supplementary Figure 3

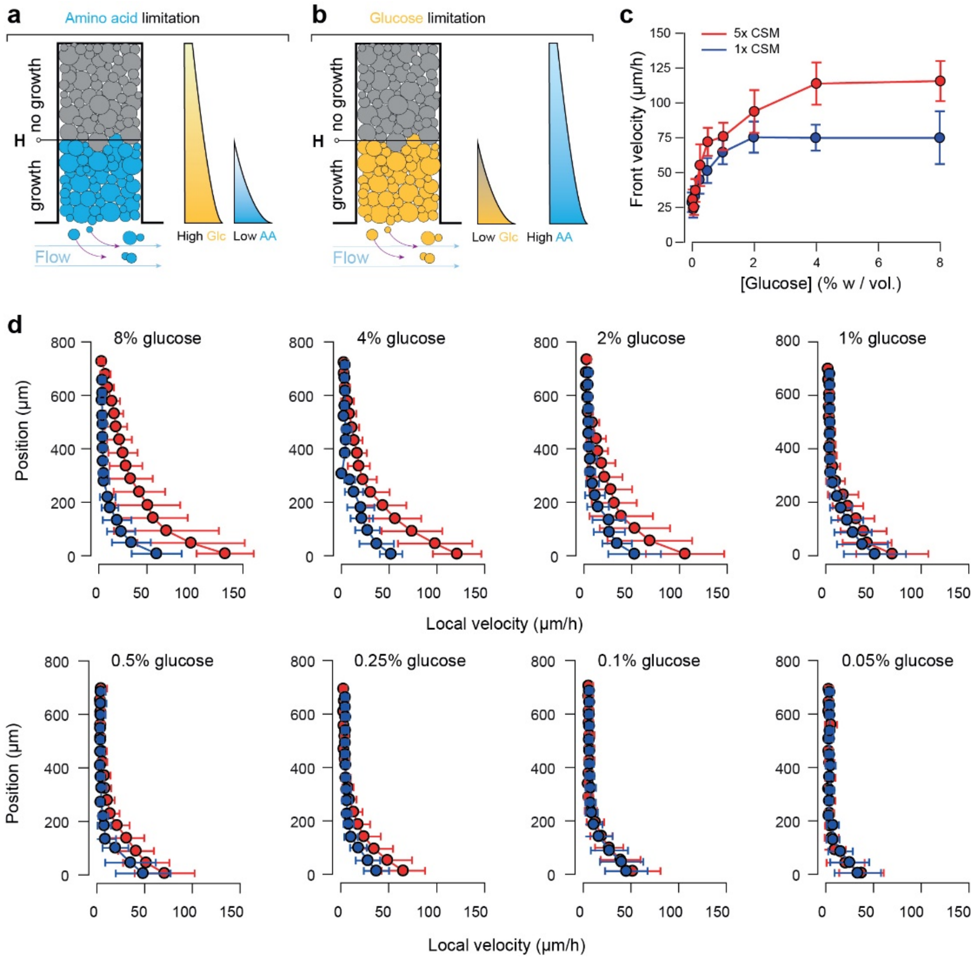

**Supplementary Figure 3. Comparison of front velocity and local velocity under low and high amino acid concentrations. S3a-b.** If not present in excess, amino acids can also limit growth of cells. Indeed, cells generate spatial gradients for all nutrients and metabolites in their microenvironment. The tradeoff between diffusion of a molecule and its uptake rate defines the properties of the gradient, thus gradients are expected to vary from one nutrient to another. In this setting, glucose and amino acids are important nutrients that can limit growth if present at too low concentrations. Glucose is required as a carbon source and the media must be supplemented with amino acids as the strain is an auxotroph. To determine the importance of amino acid concentration, we reduced the amino acid concentration to the standard concentration of SC media (called 1× CSM in the main text). **S3c**. The front velocity was lower at high glucose and low amino acid concentrations than in the case with high glucose and high amino acid concentrations. There was no significant difference up to about 1% *w/vol* (55.5 mM) glucose. Above that glucose concentration, front velocities increased with the glucose concentration at a high amino acid concentration (5× CSM), whereas front velocity levels off at a low amino acid concentration (1× CSM). Data comes from the bin closest to position 0 µm as measured in Figure 2c for each glucose concentration. Depending on the glucose concentration, minimum 5 to maximum 12 trajectories were measured (n=5-12). Represented are means and standard deviations that contain ~15-~30 velocity points each. **S3d.** The same effect was observed for local velocities. There was almost no distinction between local velocities as the source concentration of glucose increased up to about 1% *w/vol* (55.5 mM), but the difference in local velocities was striking at higher glucose concentrations. **Taken together, this means that glucose is the limiting nutrient in both 1× and 5× amino acid media, up to a glucose concentration of about 1%** ***w/vol*** **(55.5 mM).** For the case of 5× amino acid concentration, glucose remains the limiting factor for glucose concentrations used in this study. Data comes from >100 trajectories, depending on different glucose concentration. Velocity datapoints were then binned into 16 equally spaced position points. Represented are means and standard deviations of each bin.

### Supplementary Figure 4

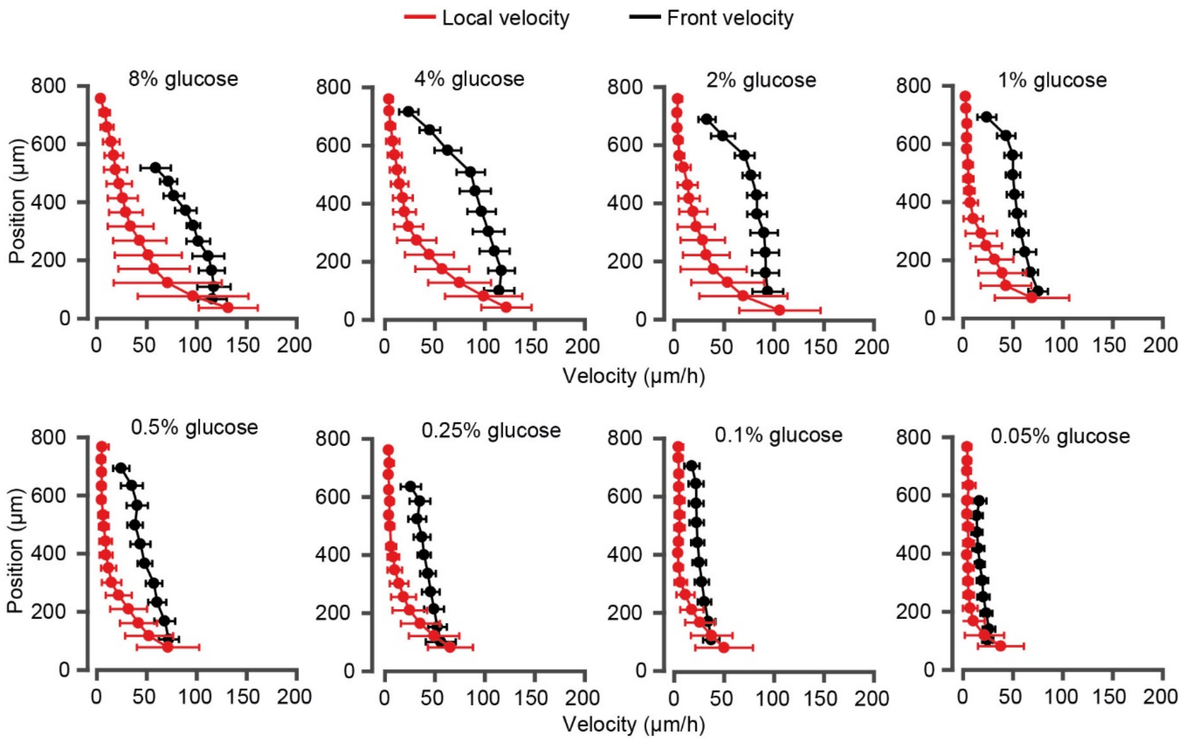

**Supplementary Figure 4. Local velocity and front velocity over a range of external glucose concentrations, $C_0$.** Red data represent the local velocity field as a function of position inside a fully developed monolayer. Error bars are +/- standard deviations computed for numerous single-cell trajectories (> 100). Black data are the front velocity measured for an expanding yeast monolayer. Maximum front velocities are reported as function of glucose concentration in Figure 2. The maximum velocities for both measurements lie within the error bars.

### Supplementary Figure 5

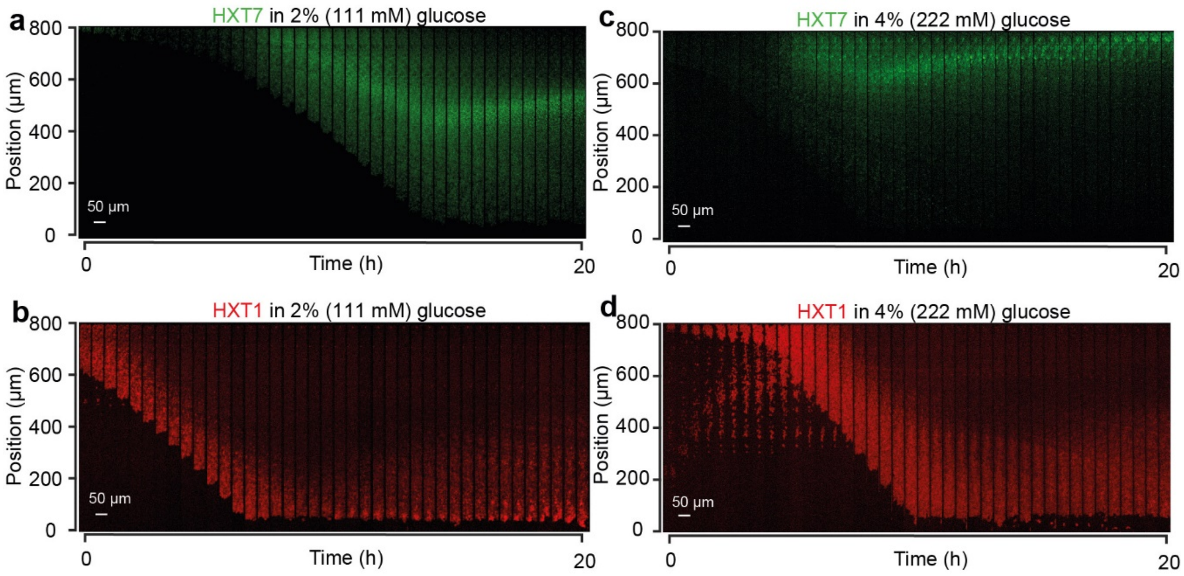

**Supplementary Figure 5: HXT1 and HXT7 landscape dynamics. S5a.** Expansion of a monolayer of cells expressing HXT7-GFP with a boundary glucose concentration of 2% *w/vol* and 5× CSM. One can observe the appearance of a peak of expression within sufficiently long monolayers. After some transient changes, the peak of expression reached steady-state position. **S5b**. Same experiment but with HXT1-GFP (shown in red here and in the main article). The landscape of HXT1 evolved very slowly over time, probably as the result of slow changes in cellular physiology and aging of the monolayer. All landscape profiles shown in the main article were measured after 10 h of growth as a fully developed monolayer. **S5c**. HXT7-GFP landscape obtained for a 4% *w/vol* boundary glucose condition. The peak of expression is very close to the end of the chamber. **S5d**. HXT1-GFP landscape obtained for a 4% *w/vol* boundary glucose condition. HXT1 landscape has a higher intensity and a deeper penetration than in **S5b**.

### Supplementary Figure 6

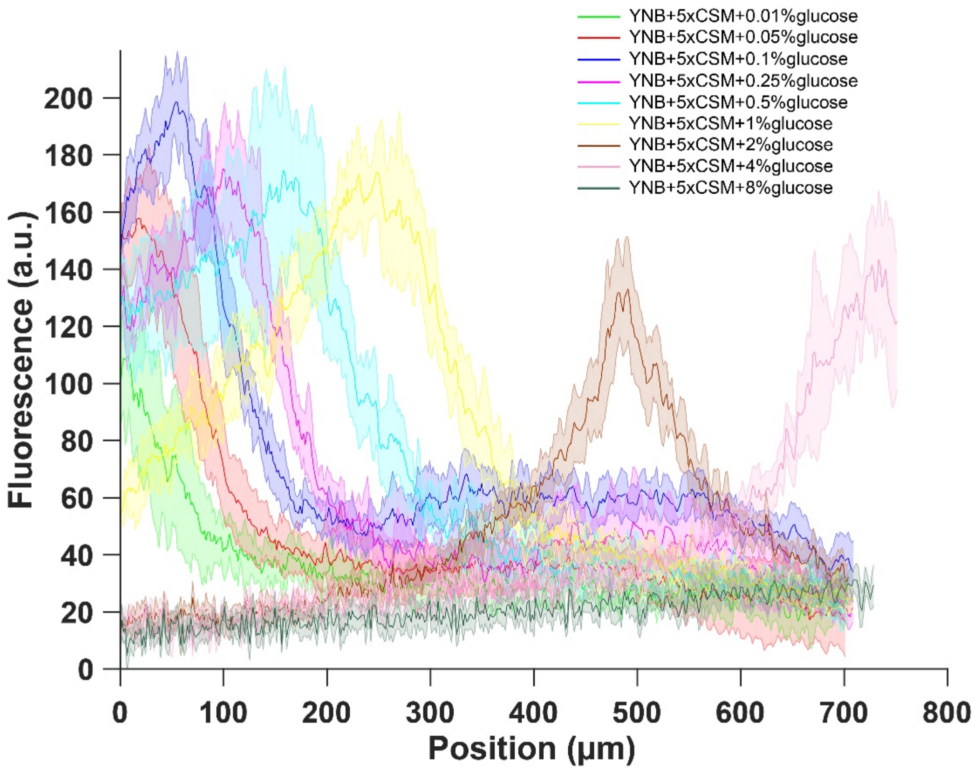

**Supplementary Figure 6. Extended figure of Figure 3c.** Gene expression of glucose transporter HXT7 in different glucose concentrations and high amino acid concentration (5x CSM). Data obtained from n=8-17 replicates per glucose concentration. Shaded error bars denote +/- one standard deviation.

### Supplementary Figure 7

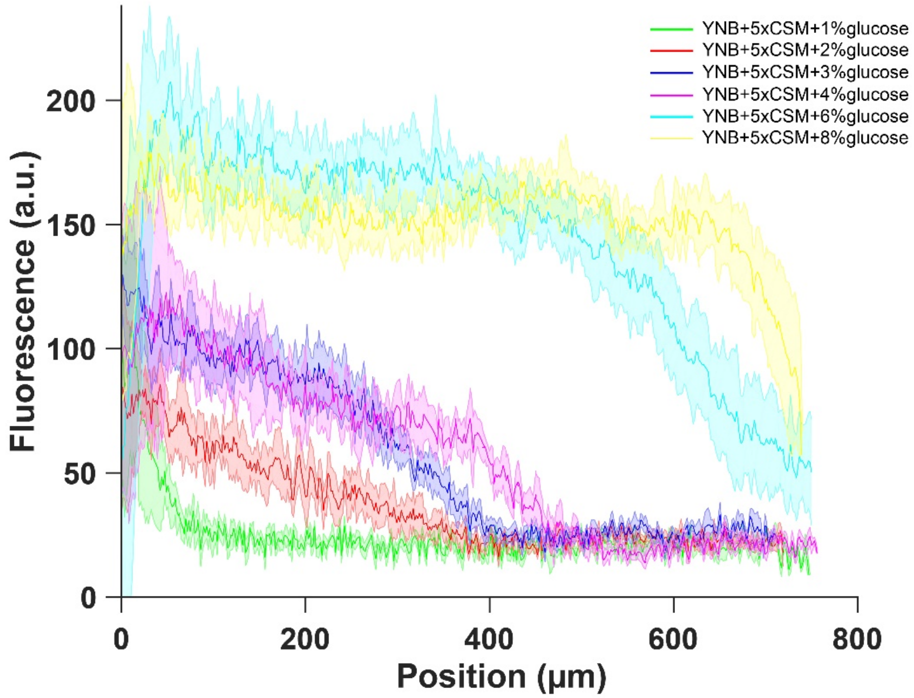

**Supplementary Figure 7. Extended figure of Figure 3e.** Gene expression of glucose transporter HXT1 in different glucose concentrations and high amino acid concentration (5x CSM). Data obtained from n=8-9 replicates per glucose concentration. Shaded error bars denote +/- one standard deviation.

### Supplementary Figure 8

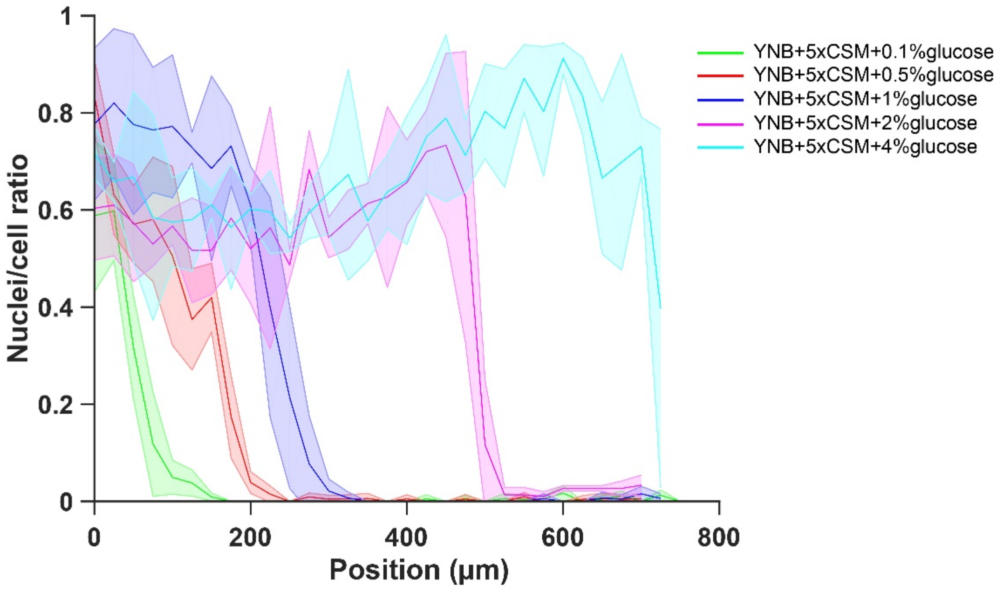

**Supplementary Figure 8. Extended figure of Figure 5b.** Gene expression of transcription factor MIG1 in different glucose concentrations and high amino acid concentration (5x CSM). Data obtained from n=3 replicates per glucose concentration. Shaded error bars denote +/- one standard deviation.

### Supplementary Figure 9

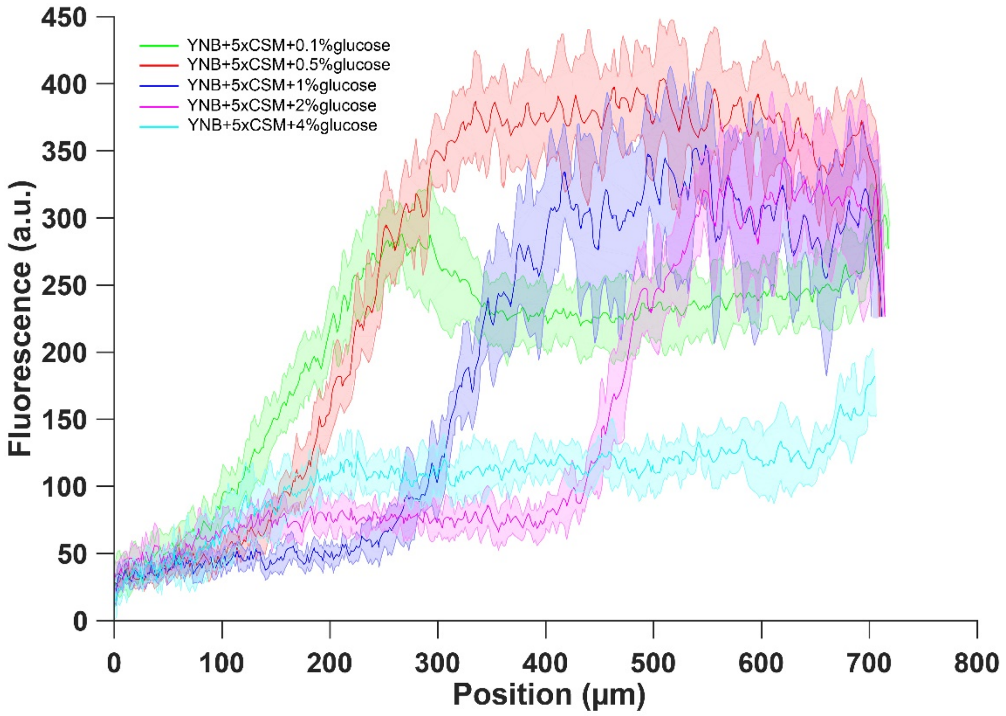

**Supplementary Figure 9. Extended figure of Figure 5a.** Gene expression of glucose transporter HXT5 in different glucose concentrations and high amino acid concentration (5x CSM). Data obtained from n=9-17 replicates per glucose concentration. Shaded error bars denote +/- one standard deviation.

### Supplementary Figure 10

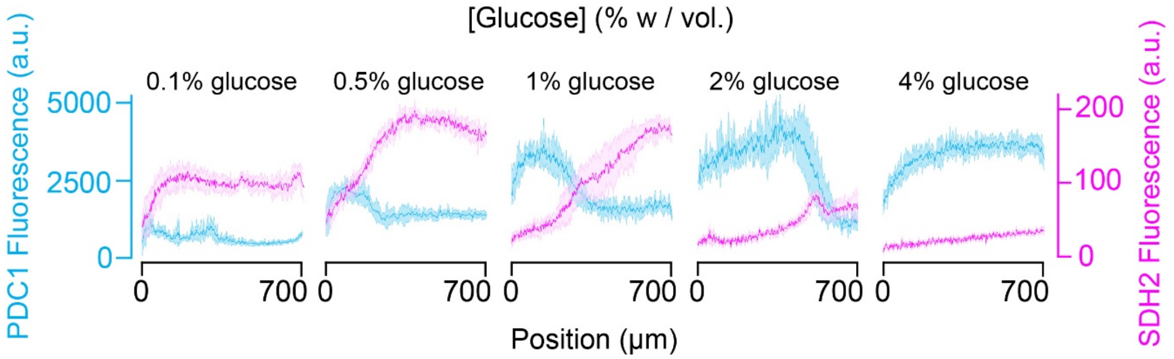

**Supplementary Figure 10. Extended figure of Figure 6a.** Gene expression landscape of pyruvate decarboxylase PDC1 (blue) and succinate dehydrogenase SDH2 (pink) in different glucose concentrations and high amino acid concentration (5x CSM). PDC1 is overexpressed during fermentations and SDH2 is overexpressed during respiration. Position 0 µm is the position of the glucose source. In colonies, two subpopulations differentiate. One that has fermentative metabolism where glucose is abundant and the other that has respiratory metabolism where glucose is scarce. Depending on the starting glucose concentrations, one or the other dominates. Data obtained from n=6-9 replicates per glucose concentration. Shaded error bars denote +/- one standard deviation.

### Supplementary Figure 11

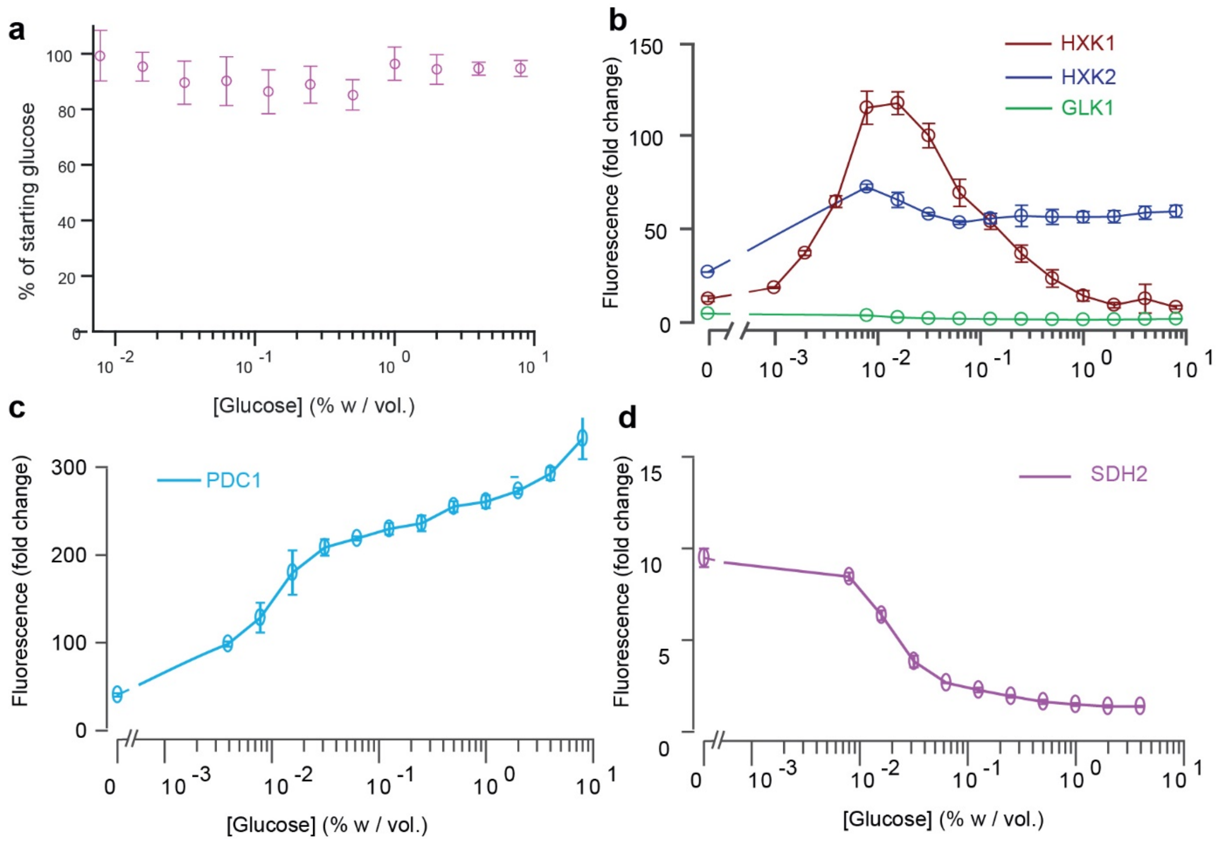

**Supplementary Figure 11.** Fluorescence intensity of key glucose metabolism genes measured by FACS over a range of glucose concentrations in batch culture. **S11a.** We also measured the glucose concentrations of the samples that were assessed by FACS. In all experiments, the glucose concentration was within 90-100% of the initial glucose concentration, indicating the cells did not use large amounts of the glucose supplied. This sets a typical error of about 10% on the inferences of glucose concentrations and gradient reconstruction in our article. Data comes from 3 replicates (n=3). **S11b.** Level of expression of HXK1 (red), HXK2 (blue) and GLK1 (green) reported in the main article represented in log-scale. **S11c.** Level of expression of PDC1 reported in the main article represented in log-scale. **S11d.** Level of expression of SDH2 reported in the main article represented in log-scale. Data in S11b, c, d comes from 3 to 6 replicates, depending on the glucose concentration (n=3-6).
